# intern: Integrated Toolkit for Extensible and Reproducible Neuroscience

**DOI:** 10.1101/2020.05.15.098707

**Authors:** Jordan Matelsky, Luis Rodriguez, Daniel Xenes, Timothy Gion, Robert Hider, Brock Wester, William Gray-Roncal

## Abstract

As neuroscience datasets continue to grow in size, the complexity of data analyses can require a detailed understanding and implementation of systems computer science for storage, access, processing, and sharing. Currently, several general data standards (e.g., Zarr, HDF5, precompute, tensorstore) and purpose-built ecosystems (e.g., BossDB, CloudVolume, DVID, and Knossos) exist. Each of these systems has advantages and limitations and is most appropriate for different use cases. Using datasets that don’t fit into RAM in this heterogeneous environment is challenging, and significant barriers exist to leverage underlying research investments. In this manuscript, we outline our perspective for how to approach this challenge through the use of community provided, standardized interfaces that unify various computational backends and abstract computer science challenges from the scientist. We introduce desirable design patterns and our reference implementation called `intern`.

## 1 Introduction

In response to the growing number and size of large-scale volumetric neuroscience datasets, the community has developed a diverse set of tools and storage frameworks that balance ease of data manipulation and storage with efficiency and cost. These tools are often purpose-built, and feature team- or task-specific features that make them particularly well-suited for their host projects, such as version control, cloud-native implementations, efficient caching, multi-tier storage, targeted annotation or proofreading tasks and more [1, 2, 3, 4, 5]. Historically, this has been advantageous, as it has enabled teams to develop tools quickly and effectively to address unique research challenges. This diverse ecosystem, however, has also led to community fragmentation and interoperability challenges because research organizations rely on standards for data storage and access that are often incompatible. As scientific questions continue to grow in ambition and scope, it is increasingly important that scientists are able to easily analyze, collaborate, and share their data using consistent formats and data-storage engines.

Though it is tempting to develop prescriptive data formats and standards, the fast-moving pace of the big-data neuroscience field — as well as the need for backward-compatibility with ongoing and past projects — will complicate the process of standardization. Instead, it is more feasible to *standardize in abstraction:* Rather than developing common data formats, it is more effective to build common data access strategies which can be applied to a variety of underlying datastores, file formats, and interfaces.

In response to collaborations that span data sizes from megabytes to petabytes, and that span institutional, international, and interdisciplinary boundaries from neuroscience to computer science to graph theory, interfacing tools are critical to reducing barriers for new and experienced scientists and enabling existing algorithms to scale to big data challenges. Data access toolkits and analysis tools (e.g., neuPrint[6], CloudVolume[7]) provide well-integrated solutions for their use cases.

We have developed *intern*, a Python client library for neuroscience data access. *intern* simplifies data transit between industry-standard data formats, and exposes a consistent and intuitive API for end-users so that code for an analysis performed on a dataset in a particular datastore format may be trivially ported to other datasets and datastores (i.e., ecosystems).

We explain our architecture and implementation details, and share several use cases common to scientific analysis which are simplified through the use of *intern*. We believe that this tool is helpful in providing seamless solutions when switching between cloud native, local, and file-based solutions, and offers an extensible software-design paradigm as new solutions are developed.

## 2 Tools

Most connectomics data management tools act as either a *data-storage* tool, which manages the (longterm) preservation of data, or a *data-access* tool – which enables an end user (whether human or automated) to access and interact with the data.

### 2.1 Data Storage Tools

Though many biological science disciplines rely on local, single-file data storage systems (e.g., HDF5, multipage TIFFs), the field of connectomics realized the need for reproducible, shareable, scalable datastores early in its evolution. These datastores are persistent, performant servers of volumetric data, and are often centralized into repositories holding information from multiple experiments and laboratories [3, 1, 4, 7, 5]. As the size of data increased, these datastores specialized in returning subvolumes of data based upon 3D user queries, rather than trying to transmit full datasets. Almost all of the most widely-used data storage tools now leverage *chunked storage* [8], an access-efficiency paradigm borrowed from domains such as astronomy and GIS [9]. This enabled databases to increase their bandwidth and serve more data-requests per second, because each subvolume could be accessed in parallel, reducing the file input/output and hard-drive read-speed bottlenecks.

Eventually, some datastores, including bossDB [1] and CloudVolume [7], moved to cloud storage systems such as Amazon AWS S3 [10] or Google Cloud Storage (GCS) [11]. These systems abstracted file-access even further and enabled high-speed network read- and write-operations, at the cost of renting — rather than owning — data storage space. While tools such as Knossos or DVID may be run on cloud resources as easily as on local compute infrastructure, other data-stores such as bossDB are cloud-native, meaning that they fully leverage the scalability and parallelism of cloud-compute resources, and *cannot* be run on conventional compute hardware.

Data storage tools can be classified into two other large categories: Those *with* server-side compute resources, and those *without*. Tools like DVID, bossDB, and Knossos use devoted compute resources that perform functions such as mesh generation, cache management, and access-control authorization. Systems like CloudVolume or zarr-backed datastores require simpler infrastructure to run, but cannot perform processes such as skeleton- or mesh-generation without client-side compute resources.

### 2.2 Data Access Tools

Some researchers may feel comfortable accessing data directly from one of the storage tools listed above (e.g., via RESTful services or object-level access), but most prefer to interact with the data through more familiar and intuitive interfaces, such as a Python library or a web interface. Almost every data storage tool mentioned above has its own devoted data *access* tool: DVID has Go and Python libraries; data stored in the *precomputed* format may be accessed with the *cloud-volume* Python library. BossDB may be accessed with either *cloud-volume* or *intern* Python libraries. A common frustration in the connectomics community is that with only a small handful of exceptions, though the underlying data may be the same in several data storage tools, most access tools are only capable of reading from their “partner” storage tool, and the interfaces vary in complexity and format. In order to integrate data and tools from our collaborators, we expanded our initial data access tool *intern* to support more data formats as well as more data storage systems, in a *Resource*-based system. *intern’s* architecture was expanded to communicate with CloudVolume-accessible volumes, DVID-hosted datasets, and several other commonly-used data storage tools and formats.

Additionally, we believe that in order to enable cross-institutional collaboration in the community, it is important to bridge the gap between those data storage tools *with* server-side compute and those without. For this reason, we also introduced a *Service*-based system into *intern* that enables a user to run surrogates for the server-side processing tools of one data-storage system using the data from another. For example, we want a user to be able to request mesh representations of data from bossDB — a tool that supports server-side mesh generation — as well as from *cloud-volume* — a tool that supports client-side mesh generation — as well as from a dataset residing in a zarr archive in S3 — a storage technique that does not support mesh generation at all. That these three tools differ in how their meshes may be generated should not matter to an end-user: The user should be able to use the same syntax to request mesh data from all of them with minimal code changes.

Finally, we wrote *intern* to be easily extended to additional use cases and features as scientific needs grow. We believe the underlying design principles are common to many research questions and have value beyond the specific implementation described here.

**Figure. 1:**
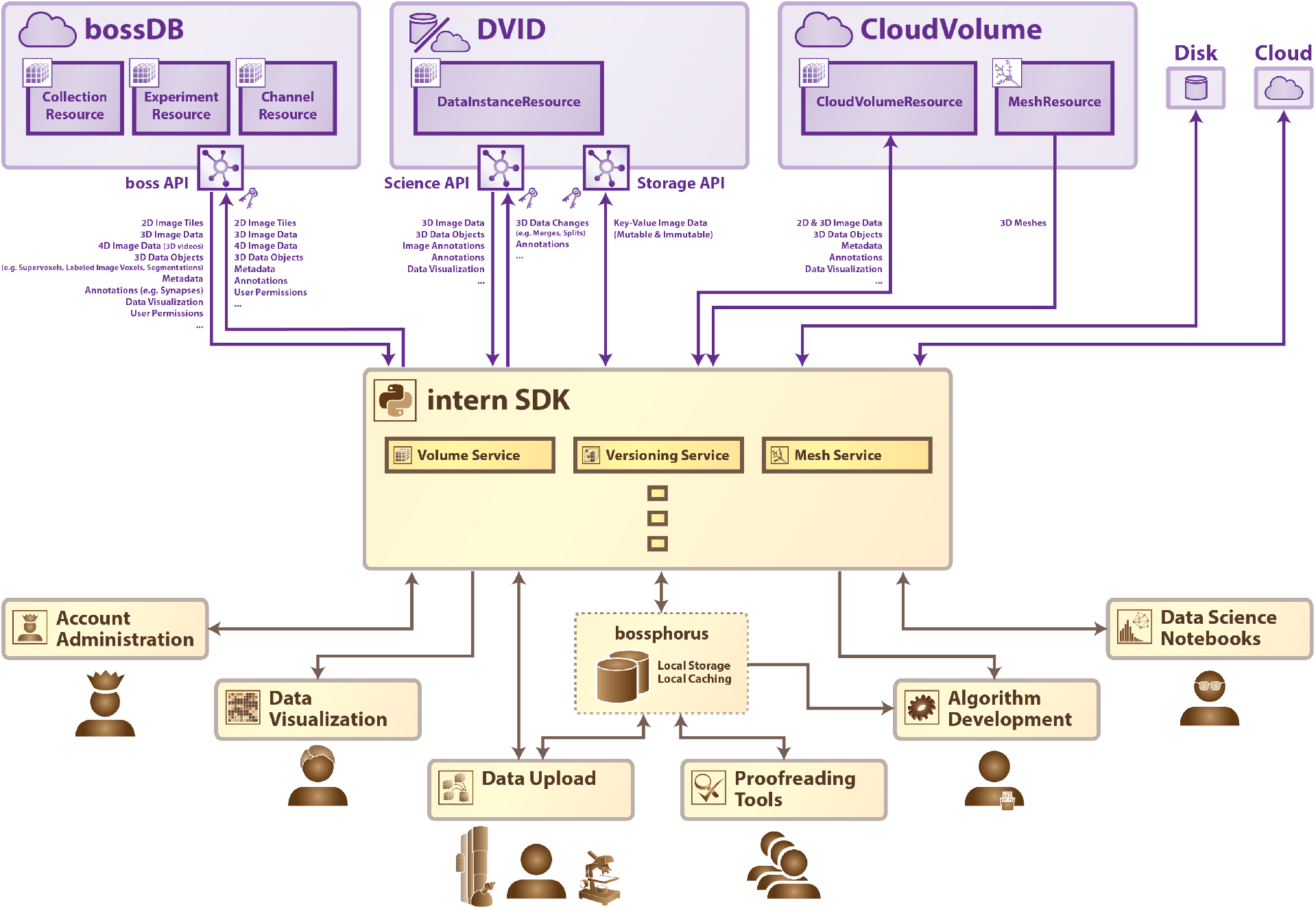
The *intern* Python library acts as a shock absorber to provide a consistent API to researchers, tool developers, and other users. A community-facing data-access tool should operate on all major data-storage systems (including CloudVolume, DVID, and bossDB), and remain flexible enough to enable common use cases (such as visualization, data upload/download, and data proofreading) without sacrificing performance.

### 2.3 The Connectomics Data-Access Ecosystem

When considering the data storage engine for a particular scientific question, several different factors should be considered, which we summarize as data size, versioning, authentication and user management, cloud services, performance, and accessibility/sharing. Each tool has a user community and powerful feature-sets: File-based solutions are simple and easily portable and understood, but are difficult to access and analyze by communities. Cloud-Volume excels in portability and simplicity, but does not provide user accounts, differential permissions, or data management services. DVID offers an excellent solution for terascale solutions and fast, efficient data-versioning, but does not leverage cloud-scale capabilities or advanced user management. BossDB is a managed cloud-native service with user permissions, access control, and a robust storage engine tested to hold and process petabytes of data, but cannot run locally and requires more infrastructural complexity than many research labs may have the expertise to maintain.

Although for the uninitiated these storage solutions may seem to introduce unnecessary complexity, managing and manipulating such large datasets and corresponding analysis derivatives (e.g., metadata) requires advanced technology. The intent of the paradigm introduced in this paper is to abstract from the user all of the challenges introduced by the scale of the data in order to allow methods to be easily run on these data while minimizing impedance mismatches.

## 3 intern

Our *intern* library implements the philosophy of abstracting computer science requirements by offering consistent data access *trait interfaces*, which are categorized into *Services, Resources*, and *Remotes*. This system of abstraction acts as a shock absorber to differing data formats, data processing, and tool functionality, and serves to enable reproducible and extensible connectomics analysis. We describe our *intern* reference implementation, and explore how other tool-developers may choose to expand *intern* or develop their own community-ready software using the same paradigm.

### 3.1 Architecture

As the field of connectomics evolves rapidly, a library must strike a balance between accessibility and adaptability. We designed our toolkit such that even minimal coding skills and copy-pasting of simple design patterns can be leveraged to reduce user burden. As the community continues to formalize use cases and data storage paradigms, programmatic workflows like SABER [12], LONI [13], Luigi [14], or other workflow managers [15, 16, 17] may allow for additional simplification and can directly leverage these functions. Point and click graphical interfaces may also follow.

In order to facilitate extension of the *intern* Python library by the community, we have published extensive online documentation for both software engineering beginners as well as professionals. The library is split into three types of trait-based interfaces; `Remotes, Resources, and Services`.

#### 3.1.1 Remotes

`Remotes` represent data storage tools, such as databases, on-disk chunked or non-chunked files, and other providers of volumetric-data access APIs. A `Remote` must at least allow the retrieval of volumetric data, and may allow upload, manipulation, user permissions, or project management as well.

#### 3.1.2 Resources

*Resources* are pointers to atomic units or groupings of data from a *Remote*. For example, in the hierarchical bossDB data paradigm, the *BossResource* implementation interfaces with a *CollectionResource*, an *ExperimentResource* and a *ChannelResource* [1]. In the *DVIDRemote* implementation, a *DataInstanceResource* points to a specific dataset at a specific version in its history.

#### 3.1.3 Services

*Services* are features or manipulations that act upon data retrieved from a *Remote*. *Service*s either call upon the server-side compute of a *Remote*, or instead a *Service* may implement a standalone local algorithm that can act as a surrogate for a *Remote* that does not have such a service available. For example, a *Cloud-VolumeRemote* has an associated *CloudVolumeMesh-Service* that invokes the built-in cloud-volume meshing functionality, but a *ZarrRemote* may use a similar, locally-executed *MeshService* with the same API.

Provided the underlying data are the same, the output from different *Service*s will be consistent (give-or-take obvious differences in performance/timing or scalability, as well as differences in parameters). In this way, raw image data from any database (i.e. *Remote*) can be treated the same; segmentation from any database can be treated the same; and annotation byproducts can be treated the same.

### 3.2 Use Cases

#### 3.2.1 Transferring data between *Remotes*

Since *Remotes* provide unique task-specific capabilities that are exclusive to a particular data store or data type, a common use-case of *intern* is to transfer data between remotes to leverage their unique capabilities.

For example, DVID provides best-in-class data-versioning of large scale image segmentation, and it may be preferable to use DVID for this sort of data-versioning rather than try to replicate this feature in other datastores. Volumetric data that is stored in, e.g., bossDB can be downloaded from the cloud for local processing and uploaded into a DVID repository using *intern*. Once the proofreading is completed, the final annotated data can be re-uploaded to bossDB in order to be cached internationally and served publicly.

#### 3.2.2 Shock-Absorption

Though such software abstractions place an additional engineering burden on developers, we assert that developing flexible, ecosystem-agnostic tools is a fundamental need of the dynamic connectomics community in lieu of more formal data-standards. To meet this requirement, we developed *intern* with such flexibility in mind: *intern* acts as a “shock-absorber” for common connectomics use-cases by implementing database-agnostic *Services* (e.g. mesh generation, skeleton generation, segmentation proofreading), which can run regardless of data source. An *intern Service* definition includes a list of its required *Resources*, and any *Remote* or other data-source that meets this interface can run the *Service*.

As a concrete example, a local marching-cubes *MeshService* converts 3D segmentation to OBJ- or *precomputed*-formatted meshes. This *Service* requires only a *VolumeResource* provider, and so it can run on, for example, a *BossRemote*, a *CloudVolumeRemote*, or even, e.g., on a raw *ZarrVolumeResource*.

Though this may appear to add unnecessary complexity, this approach enables the end-user to reproducibly run the same analysis code, changing only one line to specify from where the data should be pulled. In other words, a user may confidently change a line of code from `BossRemote#mesh(id)` to `DVIDRemote#mesh(id)`, regardless of whether the data-sources themselves support the meshing operation.

#### 3.2.3 Local Data Caching

Like many projects in the big-data neuroscience community, one of the most painful bottlenecks in much of our work is the speed with which data can be uploaded and downloaded from user-facing machines for visualization and analysis. In order to mitigate this challenge, we developed *Bossphorus*, a data relay that uses *intern* to fetch data from “upstream” data storage tools in their respective dialects and which serves data “forward” in the bossDB-flavored REST API dialect [1]. As a result, *Bossphorus* instances can be daisy-chained as a multi-tier cache. This enables an end-user to quickly browse data from a variety of sources with low latency, even if the datastore in question does not support caching. With a *Bossphorus* instance running locally using our publicly available Docker image, or a *Bossphorus* instance running on on-premise hardware at an academic institute (or indeed with both running in series), a user can interactively browse large volumes of data from multiple data sources with sub-second latency. This enables realtime data manipulation and visualization. *intern’s* interfaces are designed to be highly compatible with common data-science tools like *numpy*[18] and *pandas*[19]; popular data standards like DataJoint [17]; as well as visualization tools such as *neuroglancer*[20], *substrate*[21], *matplotlib*[22], and *plotly*[23].

#### 3.2.4 Processing

Tool and algorithm developers commonly target specific data storage ecosystems in order to reduce the burden of supporting several disparate ecosystems and data-standards. By leveraging shock-absorber tools like *intern*, algorithm developers can write code once and deploy it to a variety of datastores. As a proof of concept, we adopted a synapse-detection algorithm based upon the U-net architecture [12, 24]. This algorithm *Service* targets data downloaded from an *intern VolumeResource*, which means that it is trivially portable to data downloaded from any supported volumetric data storage service.

Just as tool designers can use *intern* to develop and test their software, the *intern* Python library is production-ready, and is verified to work at petabyte scale. We believe that reproducible and repeatable algorithm design extends past tool-design, and continues to be a fundamental aspect of responsible computational science in public-facing research. Flexible tools like *intern* equip peer institutes and collaborating researchers with the ability to quickly and accurately reproduce, verify, and build upon scientific claims.

#### 3.2.5 Visualization and Meshing

*intern*’s *Remote*, *Resource* and *Service* based architecture allows all *Remote* data-stores to benefit from all implementations of *Services*. An example of this is *intern’s MeshService*, which allows users to generate meshes using local compute resources. Any *Remote* that implements volumetric data retrieval as a *VolumeResource* (namely, all currently implemented *Remotes*) will automatically have this meshing capability. Most impactfully, due to this trait-based architecture, any future *Remote* implementation for new databases or data standards will likewise have this meshing capability without any further development required.

Any *Service* can also be used independent of the rest of the *intern* library. The *MeshService* described above, for example, will produce a *Mesh* object when passed a volume of 3D data either as an *ndarray* or as a *VolumeResource*. This mesh object can then be converted into the common obj format or into the Neuroglancer *precompute* format [20].

## 4 Discussion

In this work we highlight data accessibility, a common challenge in contemporary computational neuroscience, which has become particularly acute as data volumes grow in size and data ecosystems proliferate. New and experienced users will benefit greatly by adopting the concept of a computer science shock-absorber, which we illustrate in our solution (*intern*). Such tools are particularly valuable in domains such as connectomics, where cross-institutional collaborations and data reuse are not only common but increasingly necessary. Other complementary APIs and software libraries also exist to support approaches in the field and are well-suited for particular ecosystems and workflows. Many of these tools offer solutions that abstract many of the most challenging and repetitive aspects of large scale neuroscience discovery and also avoid common errors of interpretation. This work directly addresses the retrieval of volumetric data products but not object-level metadata such as synapse or neuron attributes, or the algorithms used to create derivative data products; these aspects are also important to consider when building standardized analysis workflows.

By developing user-facing tools such as *intern* that are flexible and provide an integrated interface to key community data storage systems, the connectomics community will be able to greatly benefit from shared, collaborative science, as well as large-scale, public, easily-accessible data.

## Author Contributions

JKM and WGR conceived of the research topic and wrote the manuscript with input from all authors. JKM, WGR, LR, DX, and TG wrote the intern library and designed the interfaces to various data stores. BW and WGR supervised the work and provided insight into community needs and usecases.

## Acknowledgments

We thank Joshua Vogelstein and Randal Burns for discussions that informed the design and vision for this project. We would also like to acknowledge Dean Kleissas for his support in the initial design and code of the intern library and the bossDB system.

## Supplemental Data

Architecture details and a user guide for the *intern* library can be found at https://bossdb.org/tools/intern.

## Data Availability Statement

The datasets analyzed for this study can be found at bossdb.org. The code used to demonstrate intern can be found at bossdb.org/tools/intern.

